# DNA-PKcs-mediated phosphorylation of AMPKγ1 regulates lysosomal AMPK activation by LKB1

**DOI:** 10.1101/409508

**Authors:** Pietri Puustinen, Anne Keldsbo, Elisabeth Corcelle-Termeau, Kevin Ngoei, Stine L. Sønder, Thomas Farkas, Klaus Kaae Andersen, Jon S. Oakhill, Marja Jäättelä

## Abstract

Autophagy is a central component of the cytoprotective cellular stress response. To enlighten stress-induced autophagy signaling, we screened a human kinome siRNA library for regulators of autophagic flux in MCF7 human breast carcinoma cells and identified the catalytic subunit of DNA-dependent protein kinase (DNA-PKcs) as a positive regulator of basal and DNA damage-induced autophagy. Analysis of autophagy-regulating signaling cascades placed DNA-PKcs upstream of the AMP-dependent protein kinase (AMPK) and ULK1 kinase. In normal culture conditions, DNA-PKcs interacted with AMPK and phosphorylated its nucleotide-sensing γγ1 subunit at Ser-192 and Thr-284, both events being significantly reduced upon AMPK activation. Alanine substitutions of DNA-PKcs phosphorylation sites in AMPKγγ1 reduced AMPK activation without affecting its nucleotide sensing capacity. Instead, the disturbance of DNA-PKcs-mediated phosphorylation of AMPKγγ inhibited the lysosomal localization of the AMPK complex and its starvation-induced association with LKB1. Taken together, our data suggest that DNA-PKcs-mediated phosphorylation of AMPKγγ primes AMPK complex to the lysosomal activation by LKB1 thereby linking DNA damage response to autophagy and cellular metabolism.

## INTRODUCTION

Macroautophagy (hereafter referred to as autophagy) is a central component of the integrated stress response in all eukaryotic cells (He & Klionsky, 2009, Kroemer, Marino et al., 2010, Weidberg, Shvets et al., 2011). It ensures the degradation of damaged or obsolete organelles, long-lived macromolecules and protein aggregates, thereby promoting the survival of starved and stressed cells. During autophagy, an isolation membrane engulfs cargo by forming a double membrane vesicle called the autophagosome, which then fuses with a lysosome to form an autolysosome where the engulfed material is degraded and recycled (Bento, Renna et al., 2016, Feng, He et al., 2014). Autophagy initiation is under tight control of at least four multiprotein kinase complexes;; the AMP-activated protein kinase (AMPK) complex, mechanistic target of rapamycin complex 1 (mTORC1), Unc-51-like kinase 1 (ULK1) complex and class 3 phosphatidylinositol 3-kinase (PI3K-III) – Beclin 1 complex (Wong, Puente et al., 2013). The AMPK and mTORC1 complexes sense the energy and nutrient status of the cell and dictate whether the cell will favor catabolic (*e.g.* autophagy) or anabolic (*e.g.* macromolecule synthesis) processes, respectively (Wong et al., 2013). Signals from AMPK and mTORC1 are integrated by the ULK1 complex that forwards the autophagy activating signals by phosphorylating a number of downstream targets, while limiting the over-activation of the pathway by suppressing the AMPK activity (Wong et al., 2013). In parallel, PI3K-III-induced formation of phosphoinositol-3-phosphate results in the recruitment of proteins that assist autophagosome formation. While the composition of the autophagy-regulating kinase complexes is emerging, our understanding of the reciprocal control of their activity and cross-talk with other cellular pathways is, as yet, limited. Recent results suggest, however, that much of this regulation takes place at the surface of the lysosome and is regulated by the availability of amino acids and glucose (Colaco & Jaattela, 2017, Dite, Ling et al., 2017, Lin & Hardie, 2018, Zhang, Jiang et al., 2014).

A wide range of DNA damaging insults continuously challenge the genome integrity. Such insults include errors encountered during DNA replication as well as physical and chemical stresses that damage the DNA directly (*e.g.* irradiation and DNA damaging chemotherapeutic drugs) or indirectly via increased production of reactive oxygen species (*e.g.* nutrient starvation and hypoxia) (Hoeijmakers, 2001). To counteract the deleterious consequences of genotoxic stress, all organisms have evolved a network of genome surveillance mechanisms designed to maintain the genomic integrity or to eliminate hazardous cells when DNA damage is beyond repair (Ciccia & Elledge, 2010, Jackson & Bartek, 2009, Okada & Mak, 2004). Recent discoveries have identified autophagy as an integrated part of the genome surveillance network that helps cells to cope with DNA damage, possibly by regulating the turnover of key proteins of DNA damage response or by supplying dNTPs essential for DNA synthesis during repair (Zhang, Shang et al., 2015). Ataxia telangiectasia mutated (ATM), the major apical kinase in homologous recombination repair of double-strand DNA breaks, is one of the putative links in this crosstalk. It can enhance autophagy by activating an AMPK activating liver kinase B1 (LKB1;; also known as serine/threonine kinase STK11) or by stabilizing the tumor protein 53 (TP53), which in turn enhances the expression of several autophagy-related genes (ATGs) and their positive regulators while suppressing the expression of autophagy-inhibiting proteins (Lee, Budanov et al., 2013, Sica, Galluzzi et al., 2015, Zhang et al., 2015). Contrary to ATM, the highly related catalytic subunit of DNA-dependent protein kinase (DNA-PKcs), which is the major apical kinase in DNA repair by non-homologous end joining (NHEJ), has been identified as a negative regulator of radiation-induced autophagy (Daido, Yamamoto et al., 2005).

In spite of the intense research in the cross talk between autophagy signaling and the genome surveillance network, several questions remain open. In order to enlighten how cells transduce the nuclear DNA damage signal to the autophagy machinery in the cytosol, we screened a human genome-wide kinase siRNA library targeting 710 known and putative kinases for regulators of DNA damage induced autophagy employing a *Renilla* luciferase (RLuc)-based reporter assay for autophagy-specific turnover of microtubule-associated protein 1 light chain 3B (LC3) in untreated and etoposide-treated MCF7 breast cancer cells (Farkas, HØyer-Hansen et al., 2009). Among several candidates, *PRKDC* siRNA encoding for DNA-PKcs emerged as the statistically strongest inhibitor of etoposide-induced autophagy as well as a statistically significant inhibitor of constitutive autophagy. DNA-PKcs being a well-characterized mediator of DNA damage response, we focused our further studies on molecular mechanisms by which it engages the autophagy machinery. Employing co-immunoprecipitation and *in vitro* kinase assays, we identified the AMP sensing *γγ*-subunit of AMPK as a DNA-PKcs substrate, and its predicted DNA-PKcs phosphorylation sites as essential regulators of the activation of the AMPK complex by liver kinase 1 (LKB1) on the lysosomal surface. These data reveal a new cytosolic function for the DNA-PKcs in the control of cellular energy metabolism.

## RESULTS

### siRNA screen for kinases that regulate DNA damage-induced autophagic flux

In order to find optimal conditions to screen the siRNA kinome library for regulators of DNA damage-induced autophagy, we performed a mini-screen of clinically relevant DNA-damaging treatments for their ability to activate autophagy in MCF breast cancer cells without inducing cell death. Autophagic flux was analyzed by a 15 h live cell assay based on a pair of MCF7 cell lines stably expressing *Renilla* Luciferase (Rluc) fused to wild type LC3B (Rluc-LC^WT^), which is degraded by autophagy, or to LC3^G^120^A^ (Rluc-LC3^G^120^A^), which is not specifically degraded by autophagy (Farkas et al., 2009). DNA damage induced by ionizing radiation (IR), etoposide, daunorubicin, doxorubicin, hydroxyurea and 5-fluorouracil (5-FU) was associated with a significant decrease in the Rluc-LC3^WT^/Rluc-LC3^G^120^A^ ratio reflecting increased autophagic flux (Fig EV1A). Etoposide was chosen as the autophagy inducer for the screen based on its potent and reproducible ability to induce autophagic flux (Fig 1A).

**Figure 1:**
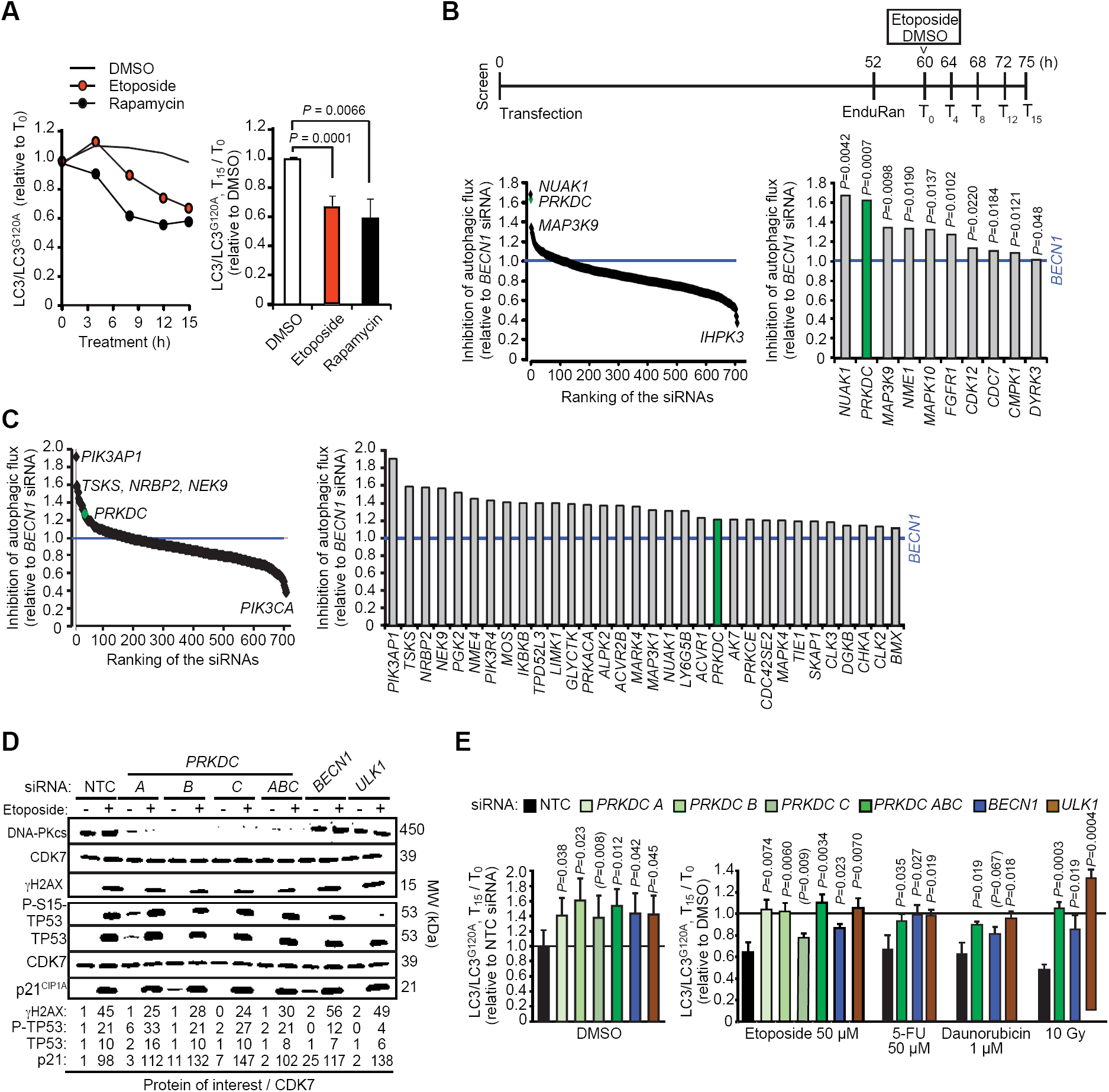
siRNA screens for kinases that regulate basal and etoposide-induced autophagy. (A) MCF7-RLuc-LC3 and MCF-RLuc-LC3^G^120^A^ cells were treated in parallel with vehicle (DMSO), 50 µM etoposide or 10 nM rapamycin (positive control) for 15 h. *Left*, kinetics of autophagic flux (reduction of LC3/LC3^G^120^A^ relative to T0) in one representative experiment. *Right*, mean ratios +SD of (LC3/LC3^G^120^A^ at T_15_)/ (LC3/LC3^G^120^A^ at T_0_) of three independent triplicate experiments. (B and C) MCF7-RLuc-LC3 and –LC3^G^120^A^ cells transfected with siRNA pools targeting 710 human kinases, *BECN1* siRNA (positive control) or non-targeting control siRNA (NTC) were treated in triplicate with 50 µM etoposide (B) or DMSO (C) for 15 h and analyzed for autophagic flux (reduction of LC3/LC3^G^120^A^) as outlined in the timeline (B, *top*). *R*anking of all siRNAs (*left*) and siRNAs with statistically significantly stronger effect than *BECN1* siRNA (*right*) are shown. (D) Representative immunoblots of indicated DNA damage response proteins in lysates of MCF7 cell transfected with indicated siRNAs for 75 h and treated with DMSO (-) or 50 µM etoposide (+) for the last 15 h. Cyclin dependent kinase 7 (CDK7) served as a loading control. (E)MCF7-RLuc-LC3 and –LC3^G^120^A^ cells were transfected with indicated siRNAs, treated as indicated for 15 h and analyzed for autophagic flux as in (B). Values represent means +SD of ≥ 3 independent triplicate experiments. P-values were calculated by 2-tailed, homoscedastic student’s *t*-test (A, C and E) or by a linear regression model and Wald test of autophagy inhibition over time (B).

We then screened a genome-wide kinome siRNA library targeting 710 known and putative human kinases or kinase-related genes (3 siRNAs/gene à 6 nM) in parallel with non-targeting control (NTC, negative control) and *BECN1* (positive control) siRNAs (18 nM) according to the timeline outlined in Fig 1B. Seventy-four siRNAs inhibited etoposide-induced autophagy to the same or greater extent than *BECN1* siRNA (Table EV1), and in ten cases the inhibition was significantly stronger than by *BECN1* siRNA when comparing inhibition kinetics (inhibition per time unit) and tested by means of Wald tests (Fig 1B and Table EV1). In normal growth conditions, 32 siRNAs inhibited autophagy significantly better than *BECN1* siRNA (Fig 1C).

*PRKDC* siRNA, which targets DNA-PKcs, emerged as the statistically most significant inhibitor of etoposide-induced autophagy with 1.6-fold stronger inhibitory effect than *BECN1* siRNA (Fig 1B;; Table EV1). It also inhibited basal autophagy significantly better than *BECN1* siRNA (Fig 1C). DNA-PKcs, which plays an essential role in DNA repair by NHEJ, belongs to the PI3K-related kinase (PIKK) family, whose other members include two other well characterized DNA damage response kinases, ATM and ataxia-and Rad3-related (ATR), as well as autophagy-regulating mTOR (Abraham, 2004). Notably, *ATR* and *ATM* siRNAs were also among the candidate inhibitors of the etoposide-induced autophagy in our screen (Table EV1).

### Depletion of DNA-PKcs inhibits basal and stress-induced autophagic flux

Prompted by its strong effect on etoposide-induced autophagy, known involvement in DNA damage signaling pathways, and relatively high mRNA expression in MCF7 cells as compared to *ATM* and *ATR* (Fig EV1B), we focused our further studies on DNA-PKcs. First, we validated its role in autophagy by transfecting MCF7-RLuc-LC3^WT^/^G^120^A^ reporter cells with three individual *PRKDC* siRNAs (6 nM), all of which effectively decreased DNA-PKcs protein levels and DNA-PKcs activity as judged by reduced etoposide-induced phosphorylation of its substrate, H2A histone family member X (H2AX) (Fig 1D). Two out of three individual siRNAs inhibited both the basal and etoposide-induced autophagic flux statistically significantly, while the third siRNA showed a similar inhibitory tendency without reaching statistical significance (Fig 1E). In addition to etoposide-induced autophagic flux, *PRKDC* siRNA pool inhibited that induced by 5-FU, daunorubicin and ionizing radiation in MCF7-RLuc-LC3^WT^/^G^120^A^ reporter cells (Fig 1E). In U2OS osteosarcoma RLuc-LC3^WT^/^G^120^A^ reporter cells, *PRKDC* siRNA pool had similar statistically significant inhibitory activity towards both constitutive and DNA damage-induced autophagy, whereas in HeLa cervix carcinoma RLuc-LC3^WT^/^G^120^A^ reporter cells, it significantly inhibited the constitutive autophagic flux and had an inhibitory tendency against etoposide-induced autophagy (Fig EV1C).

Due to the essential role of DNA-PKcs in the DNA damage response, the autophagy inhibition observed upon its depletion may be caused by defects in DNA repair. *PRKDC* siRNA affected, however, neither the etoposide-induced cell cycle arrest (Fig EV1D) nor the activation of TP53 pathway as analyzed by the level and phosphorylation of TP53 and the level of the transcriptional target of TP53, cyclin-dependent kinase inhibitor 1A (p21^CIP^1^A^) (Fig 1D).Furthermore, *XRCC5* siRNA targeting KU80 regulatory subunit of the DNAPK complex, which is essential for the function of DNA-PKcs in NHEJ (Lees-Miller & Meek, 2003), had no effect on etoposide-induced autophagic flux (Fig EV1E). These data suggest that DNA-PKcs regulates autophagy in a manner independent of its role in the DNA damage repair.

### DNA-PKcs regulates AMPK-ULK1 signaling pathway

Having established DNA-PKcs as a regulator of autophagic flux, we sought to uncover the underlying molecular mechanism. DNA-PKcs depletion inhibited etoposide-induced formation of WD repeat domain phosphoinositide-interacting protein 2 (WIPI2)-and LC3-positive autophagic membranes (Figs 2A, EV2A and EV2B), indicating that DNA-PKcs regulates signaling pathways leading to the autophagy initiation rather than those regulating autophagosome maturation and degradation. Thus, we investigated whether this regulation involved mTORC1, the major negative regulator of autophagy initiation, by using an mTORC1 inhibitor rapamycin to induce autophagy. The significant reduction in the abundance of rapamycin-induced autophagic membranes by *PRKDC* siRNA suggested that DNA-PKcs regulated autophagy in an mTORC1-independent manner (Figs 2A, EV2A and EV2B). Accordingly, DNA-PKcs depletion neither increased basal mTORC1 activity nor its inhibition by etoposide as analyzed by the phosphorylation status of an mTORC1 substrate, Tyr-389 in ribosomal protein S6 kinase β1 (T389-p70^S^6^K^1^^;; Fig 2B). Instead, *PRKDC* siRNA inhibited the AMPK-ULK1 pathway as shown by decreased basal and etoposide-induced phosphorylation of an ULK1 substrate, serine 318 in autophagy related 13 (S318-ATG13;; Fig 2C). The sensitivity of AMPK to the transfection stress made it difficult to study AMPK-ULK1 pathway in detail in this model system, which relies on transient siRNA transfections. Thus, we continued the study in U2OS osteosarcoma cells stably infected with *PRKDC* shRNA-encoding lentivirus. Akin to the transient depletion of *PRKDC* by siRNAs, its long-term depletion by shRNA significantly reduced etoposide-and rapamycin-induced formation of LC3-and p62-(also known as sequestesome 1) positive autophagic membranes (Fig 2D). Furthermore, *PRKDC* shRNA significantly reduced the activity of the AMPK-ULK1 pathway as evidenced by reduced phosphorylation of Tyr-172 in the activation site of the catalytic *αα*-subunit of AMPK (T172-AMPK*αα*), AMPK substrates, Ser-79 in acetyl-coenzyme A carboxylase *αα* (S79-ACC) and Thr-317 in ULK1 (T317-ULK1), and ULK1 substrate S318-ATG13 (Fig 2E). We could not detect the etoposide-induced activation of AMPK reproducibly in U2OS cells, possibly due to its transient and unsynchronized nature. Thus, we studied the effect of *PRKDC* shRNA following a faster and stronger AMPK activation by A769662, a small molecule that activates AMPK allosterically and by inhibiting the dephosphorylation of T172-AMPK*αα* (Sanders, Ali et al., 2007). *PRKDC* shRNA reduced the A769662-induced activation of AMPK as analyzed by the phosphorylation status of T172-AMPK*αα* and its substrates, S317-and S555-ULK1 (Fig 3A, *left*), Taken together, these data suggest that DNA-PKcs regulates autophagy independently of mTORC1, by enhancing the basal and stimulus-induced activity of the AMPK complex.

**Figure 2:**
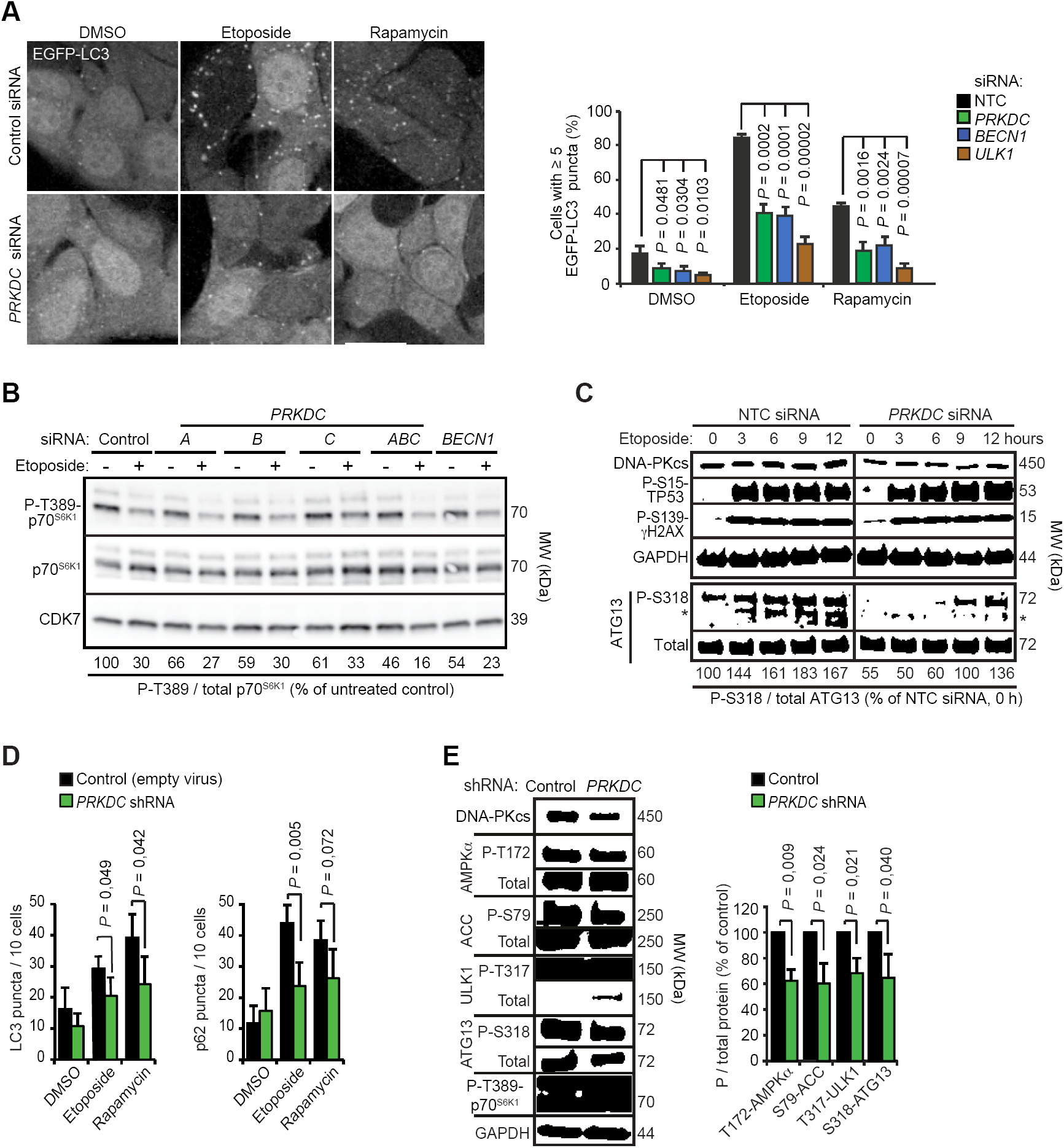
DNA-PKcs regulates autophagy in an AMPK-and ULK1-dependent manner. (A)MCF7-EGFP-LC3 cells transfected with indicated siRNAs for 75 h were treated with DMSO or 50 µM etoposide for the last 15 h or 10 nM rapamycin for the last 2 h. *Left*, representative confocal images of EGFP puncta (grey). Scale bar, 10 µM. See Fig EV2A for all images. *Right*, Mean percentages of cells with ≥ 5 EGFP-LC3 puncta +SD of three independent experiments with > 50 randomly chosen cells analyzed in each sample. (B and C) Representative immunoblots of indicated proteins in lysates of MCF7 cell transfected with indicated siRNAs for 75 h and treated with DMSO (-) or 50 µM etoposide (+) for the last 15 h or with 50 µM etoposide for indicated times (C). CDK7 (B) and GAPDH (C) served as loading controls. Values, densitometric quantifications of phospho/total protein ratios as percentages of control samples. (D)LC3B (*left*) and p62 (*right*) puncta in U2OS-control and U2OS-*PRKDC* shRNA cells treated with DMSO, 50 µM etoposide or 10 nM rapamycin for 24 h. Values represent means +SD of four independent experiments with a minimum of five randomly chosen groups of 10 cells analyzed in each sample. (E)Representative immunoblots of indicated proteins from lysates of U2OS-control and U2OS-*PRKDC* shRNA cells (*left*) and densitometric quantification of indicated phospho-protein/total protein ratios (*right*). Values represent means of three independent experiments +SD. *P*-values were calculated by 2-tailed, homoscedastic student’s *t*-test.

**Figure 3:**
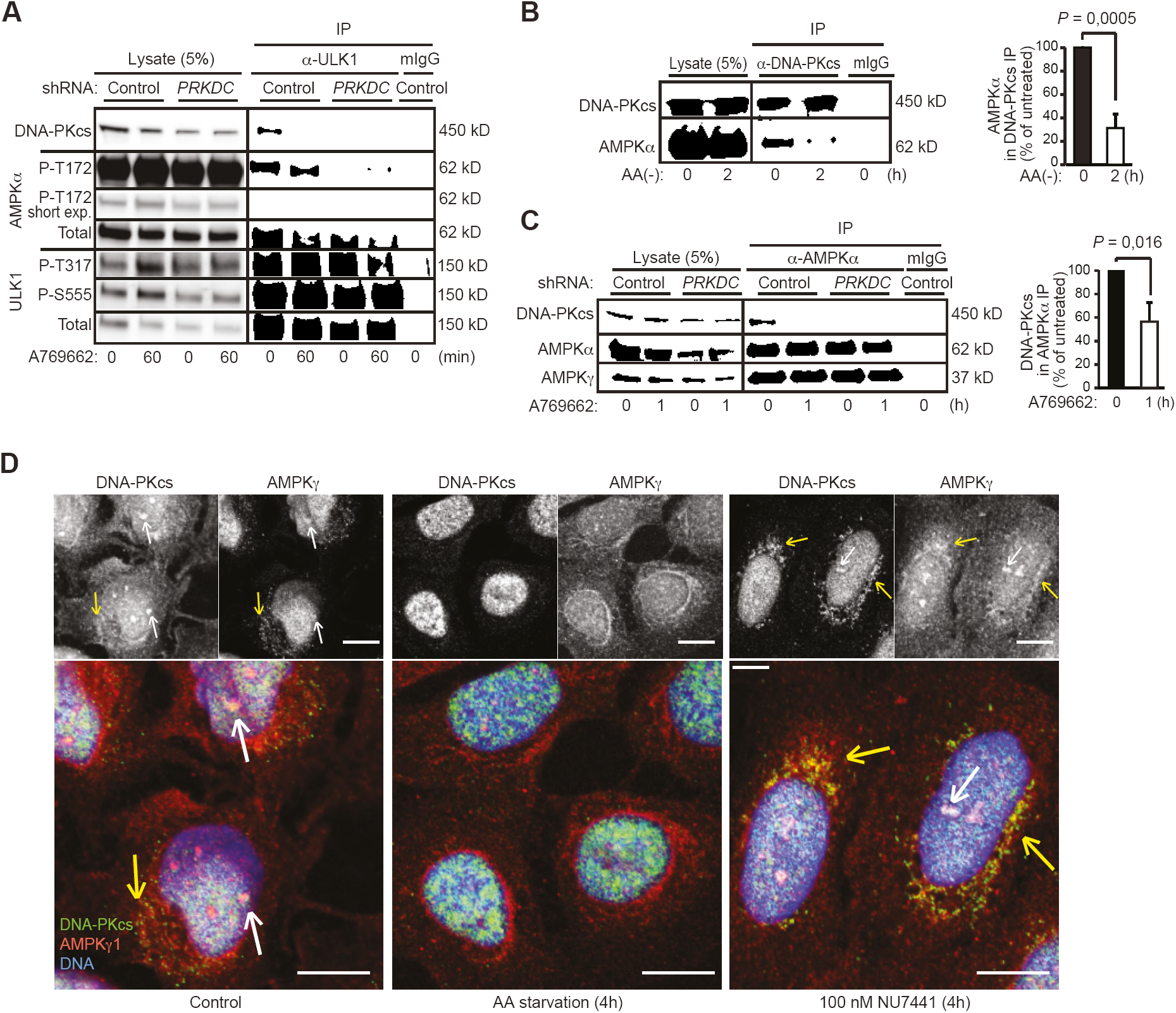
Dynamic association of DNA-PKcs, AMPK complex and ULK1. (A)Immunoprecipitation (IP) of endogenous ULK1 complexes from lysates of control and *PRKDC* shRNA-infected U2OS cells treated with DMSO (0) or 1 µM A769662 for 60 min (60). (B)IP of endogenous DNA-PKcs complexes were immunoprecipitated from lysates of U2OS cells grown in complete media (0) or starved for amino acids for 2 h (2). *Left*, representative immunoblots of lysates and IPs. *Right*, quantification of co-precipitating AMPK*αα* from three independent experiments +SD. (C) IP of endogenous AMPK*αα* complexes from lysates of control and *PRKDC* shRNA-infected U2OS cells treated with DMSO (0) or 1 µM A769662 for 1 h (1). *Left*, representative immunoblots of lysates and IPs. *Right*, quantification of co-precipitating DNA-PKcs from three independent experiments +SD. (D)Representative confocal images of MCF7 cells grown in complete medium, starved for amino acids or treated with 100 nM NU7441 for 4 h, and stained for endogenous DNA-PKcs (green), AMPK*γγ*1 (red) and DNA (Hoechst, blue). Yellow and white arrows indicate co-localization of the proteins in the cytoplasm and nucleus, respectively. Scale bar, 10 µm. *P*-values were calculated by 2-tailed, homoscedastic student’s *t*-test.

### DNA-PKcs associates with the AMPK-ULK1 complex

Prompted by the observed DNA-PKcs-dependent activity of the AMPK-ULK1 pathway, we next investigated putative associations of these kinases by immunoblot analyses of proteins co-precipitating with endogenous DNA-PKcs, AMPK complex (AMPK*αα* or AMPK*γγ*) or ULK1. In normal growth conditions, DNA-PKcs had relatively strong associations with AMPK complex and ULK1, both of which dissociated from DNA-PKcs upon AMPK activation by amino acid starvation (Figs 3B and EV3A), A769662 (Figs 3A, 3C and EV3B), glucose starvation (Fig EV3C) or etoposide (Fig S3D). Notably, the shRNA-mediated depletion of DNA-PKcs did not affect the stability of the AMPK complex (Fig 3C), but clearly reduced its association with ULK1 and AMPK-mediated phosphorylation of ULK1 as mentioned above (Fig 3A and EV3B). These data suggest that DNA-PKcs interacts with AMPK-ULK1 complex in normal growth conditions and regulates its activation by various stimuli, which eventually trigger the dissociation of DNA-PKcs from the complex. Supporting this model, immunocytochemical analyses revealed a dynamic co-localization of DNA-PKcs and AMPK complex in MCF7 cells (Fig 3D). Partial co-localization of DNA-PKcs and AMPK*γγ*observed in the cytoplasm and nuclei of untreated cells was abolished by amino acid starvation, which triggered a perinuclear accumulation of AMPK*γγ* and mainly nuclear localization of DNA-PKcs, whereas inhibition of DNA-PKcs activity by NU7441, a small molecule ATP competitive inhibitor of DNA-PK (Leahy, Golding et al., 2004), increased the co-localization of DNA-PKcs and AMPK*γγ* in the perinuclear area (Fig 3D). Taken together, our biochemical and immunocytochemical analyses suggest that a subset of DNA-PKcs associates with the AMPK complex in normal growth conditions to prime it for activation, and that the activation of the AMPK complex results in the dissociation of DNA-PKcs from the AMPK complex in a DNA-PKcs activity-dependent manner.

### DNA-PKcs phosphorylates AMPKγγ1

In order to test whether DNA-PKcs is capable of phosphorylating the AMPK complex, we performed an *in vitro* kinase assay with immuno-purified DNA-PKcs and AMPK complexes. Mixing of the two kinases resulted in an NU7441-sensitive phosphorylation of a 37 kDa protein suggesting that DNA-PKcs phosphorylates the AMPK*γ* subunit with predicted molecular weight of 37 kDa (Fig 4A). An unbiased prediction of putative phosphorylation sites in AMPK*γ*1 using Scansite motif prediction platform (http://scansite.mit.edu) provided two possible recognition motifs for the PIKK family kinases located in the third (Ser-192, predicted ATM site) and fourth (Thr-284, predicted DNA-PK and ATM site) cystathionine-b-synthase (CBS) tandem repeat motifs (Fig 4B), which act as sensors of cellular AMP:ATP and ADP:ATP ratios (Hardie, Schaffer et al., 2016). According to the available crystal structure of the holo-AMPK complex (AMPK*αα*1, AMPKβ2, AMPK*γ*1) (Li, Wang et al., 2015), the predicted phosphorylation sites are in the close proximity of each other and accessible for kinases on the surface of the protein (Fig 4B). These sites are also highly conserved in evolution from zebra fish and chicken to human and similar ATM and DNA-PKcs /ATM consensus recognition sequences were found in AMPK*γ*3, the main AMPK*γ*-subunit in skeletal muscle (Figs EV4A and EV4B). Recognition motif comparison by PhophoSitePlus^®^ online resource (Hornbeck, Zhang et al., 2015) favored, however, AMPK*γ*1 as a DNA-PKcs /ATM substrate over AMPK*γ*3. Based on this and the relatively high expression of AMPK*γ*1 in MCF7 cells (Fig EV1B), we focused our further studies on AMPK*γ*1. First, we verified the ability of DNA-PKcs to phosphorylate the predicted sites by using short peptide sequences surrounding them (PEFMSK- Ser-192-LEELQIGC and KCYLHE-Thr-284 -LETIINRLC) in an *in vitro* kinase assay. Both wild type peptides were effectively phosphorylated by DNA-PKcs, while their phosphorylated counterparts were not (Fig 4C). Moreover, an *in vitro* kinase assay using immuno-purified full length proteins as substrates showed a strong reduction in the DNA-PKcs -mediated phosphorylation of AMPK*γ*1 when Ser-192 (S192A) or Thr-284 (T284A) alone or together (AA) were mutated to alanine, confirming them as accessible DNA-PKcs substrates *in vitro* (Fig 4D). In order to study the phosphorylation of these sites in living cells, we raised phosphor-specific antibodies against them (Fig EV4C). The phosphorylation of T284-AMPK*γ*1 was clearly reduced by RNAi-mediated depletion of DNA-PKcs, pharmacological inhibition of DNA-PKcs kinase activity by NU7441, glucose starvation and etoposide treatment in MCF7 cells, whereas that of S192-AMPK*γ*1 was less affected (Figs 4E, 4F and EV4D). Akin to MCF7 cells, the phosphorylation of T284-AMPK*γ*1 but not that of S192-AMPK*γ*1, was clearly reduced in U2OS cells upon glucose starvation and DNA-PKcs depletion (Figs 4G and 4H). Supporting the role of DNA-PKcs as the principal T284-AMPK*γ*1kinase, pharmacological inhibitors of other PIKK family members, *i.e.* ATM (KU55933) or mTOR (Torin), or closely related phosphatidylinostide 3-kinases (LY294002) did not affect the phosphorylation status of T284-AMPK*γ*1 to the same extent as DNA-PKcs inhibition by NU7441 or NU7026 (Fig EV4F). Furthermore, phosphorylation of this site was not reduced by the compound c-mediated inhibition of the catalytic activity of the AMPK complex itself (Fig 4H). Notably, the reduced phosphorylation of T284-AMPK*γ*1 upon inhibition of DNA-PKcs activity was associated with reduced activation of the AMPK complex both in MCF7 and U2OS cells as analyzed by the phosphorylation status of T172-AMPK*αα* (Figs 4F and 4H). These data suggest that in cancer cells studied here, constitutive DNA-PKcs activity keeps T284-AMPK*γ*1 phosphorylated thereby promoting the activation of the AMPK complex. Indeed, the two autophosphorylation sites in DNA-PKcs, serine 2056 (S2056) and threonine 2609 (T2609), that reflect the activity of the kinase (Blackford & Jackson, 2017), were phosphorylated in MCF7 cells in optimal growth conditions, and their phosphorylation was not markedly altered by glucose starvation (Fig 4I, *left*). On the other hand, neither DNA-PKcs (S2056 and T2609) nor T284-AMPK*γ*1 was phosphorylated in primary fibroblasts whether grown in complete medium or starved for glucose (Fig 4I, *right*). The lack of DNA-PKcs-AMPK*γ*1 pathway activation in these primary cells was accompanied by the lack of AMPK complex activity (P-T172-AMPK*αα*) in optimal growth conditions and only a very weak activation upon glucose starvation (Fig 4I). Thus, the constitutive activity of DNA-PKcs in cancer cells may enhance the basal activity and stimulus-dependent activation of the AMPK complex via phosphorylation of T284 in the AMPK*γ*1 subunit.

**Figure 4:**
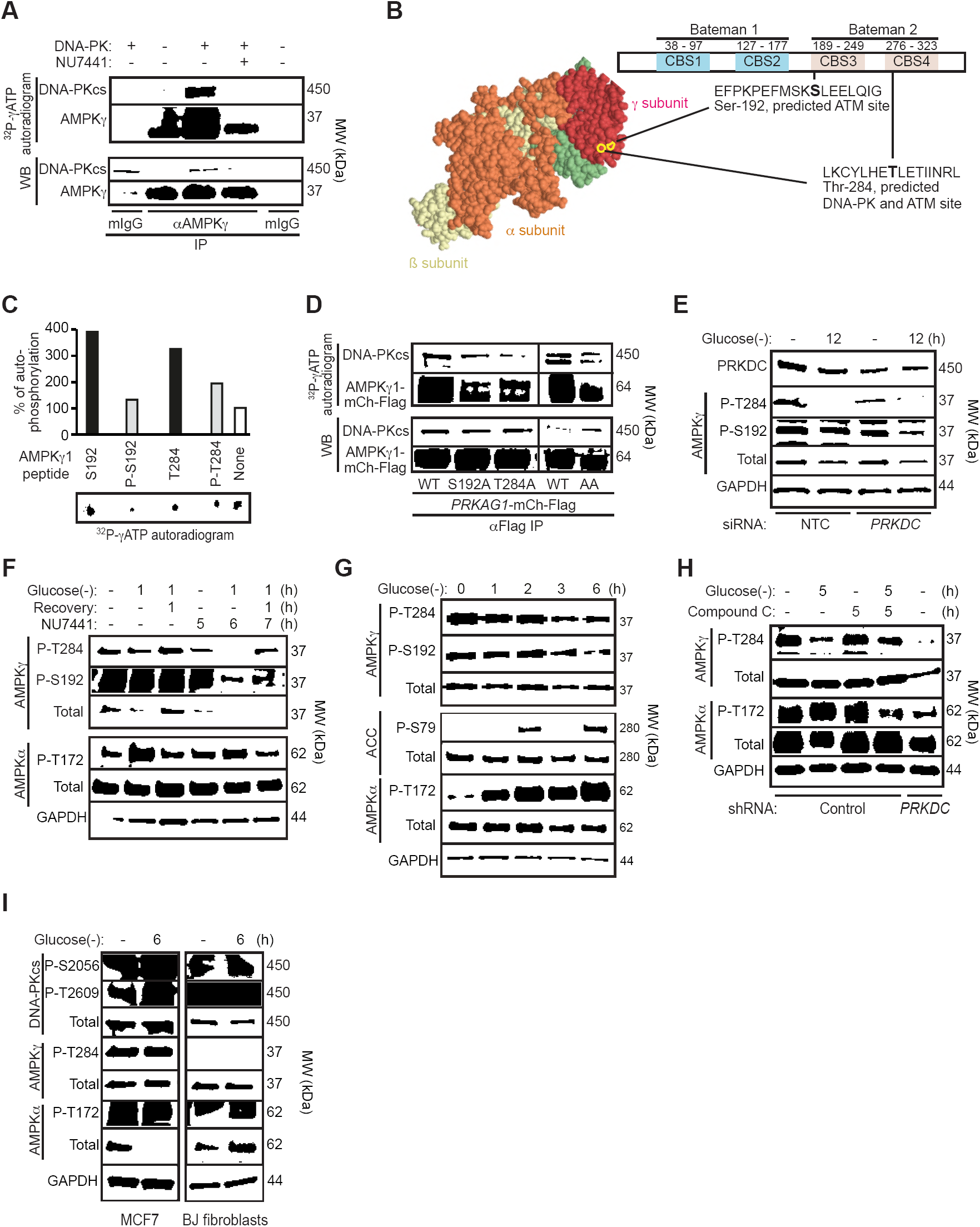
DNA-PKcs phosphorylates AMPKγ1. *(A)In vitro* DNA-PK kinase assay with recombinant DNA-PK (+) and AMPK*γ*1 immunoprecipitated from MCF7 cells in stringent conditions. When indicated 100 nM NU7441 was added to the reaction. (B)Predicted DNA-PKcs/ATM and ATM phosphorylation sites in AMPK*γ*1. *Left*, crystal structure of holo-AMPK complex consisting of AMPK*αα*1, AMPKβ2 and AMPK*γ*1 (Li et al., 2015). Ser-192 and Thr-284 are highlighted with yellow. *Right*, domain structure of AMPK*γ*1 with CBS domains 1-4 and predicted phosphorylation sites highlighted. *(C)In vitro* DNA-PK kinase assay with indicated AMPK*γ*1 peptides as substrates. Radioactivity was analyzed by FUJI phospho-imager plate and spots were quantitated with FujiFilm MultiGauge version 3.2. (D)*In vitro* DNA-PK kinase assay with wild type (WT) AMPK*γ*1-Ch-Flag or its S192A, T284A and S192A/T284A (AA) mutants immunopurified from U2OS cells as substrates. (E)Representative immunoblots of indicated proteins from MCF7 cells transfected with non-targeting control (NTC) or *PRKDC* siRNA for 60 h and starved for glucose for the last 12 h when indicated (12). (F)Representative immunoblots of indicated proteins from MCF7 cells left untreated or starved for glucose for 12 h with or without 1 h recovery. When indicated cells were retreated with 100 nM NU7441 for indicated times. (G)Representative immunoblots of indicated proteins from U2OS cells left untreated or starved for glucose for indicated times.(H)Representative immunoblots of indicated proteins from control and *PRKDC* shRNA-infected U2OS cells left untreated, starved for glucose or treated with 5 µM compound C as indicated. (I)Representative immunoblots of indicated proteins from MCF7 cells and BJ fibroblasts left untreated or starved for glucose for 6 h.

### Phosphorylation of AMPKγ1 regulates AMPK localization and activity

Having established S192 and T284 as relevant AMPK*γ*1 phosphorylation sites in cancer cells, we studied their putative role in AMPK activation and signaling. Because these sites are localized in the CBS repeats responsible for sensing intracellular AMP:ATP and ADP:ATP ratios, we first investigated whether their phosphorylation status affected the ability of AMP to activate the AMPK complex. For this purpose, we expressed heterotrimeric *αα*21*γ*1 AMPK complex consisting of GST-AMPK*αα*2, AMPKβ1-Flag and either wild type or phosphorylation-deficient mutants (S192A or T284A) of HA-AMPK*γ*1 in COS7 human kidney epithelial cell, and purified the complexes using anti-Flag affinity agarose gel. Supporting the idea that AMPK*γ*1 phosphorylation promotes AMPK activity, T172-AMPK*αα*2 was 61% and 43% more phosphorylated when in complex with the wild type AMPK*γ*1 than when in the complex with S192A and T284A mutants, respectively (Fig 5A). The mutant AMPK*γ*1 proteins formed, however, heterotrimeric AMPK complexes as efficiently as the wild type protein (Fig 5B). In an *in vitro* kinase assay, the wild type and mutant AMPK complexes had similar basal kinase activities, all of which were effectively stimulated by AMP (Fig 5B). The AMP-induced activation of the T248A-AMPK*γ*1 containing complex was similar to that of the wild type complex, whereas that of the S192A-AMPK*γ*1 containing complex was even more efficient than the wild type complex (Fig 5B). Thus, the phosphorylation of AMPK*γ*1 at S192 and T284 promotes the AMPK activity in cells without enhancing the formation of heterotrimeric AMPK complexes or the ability of AMP to bind AMPK*γ*1 and activate the AMPK complex.

**Figure 5.**
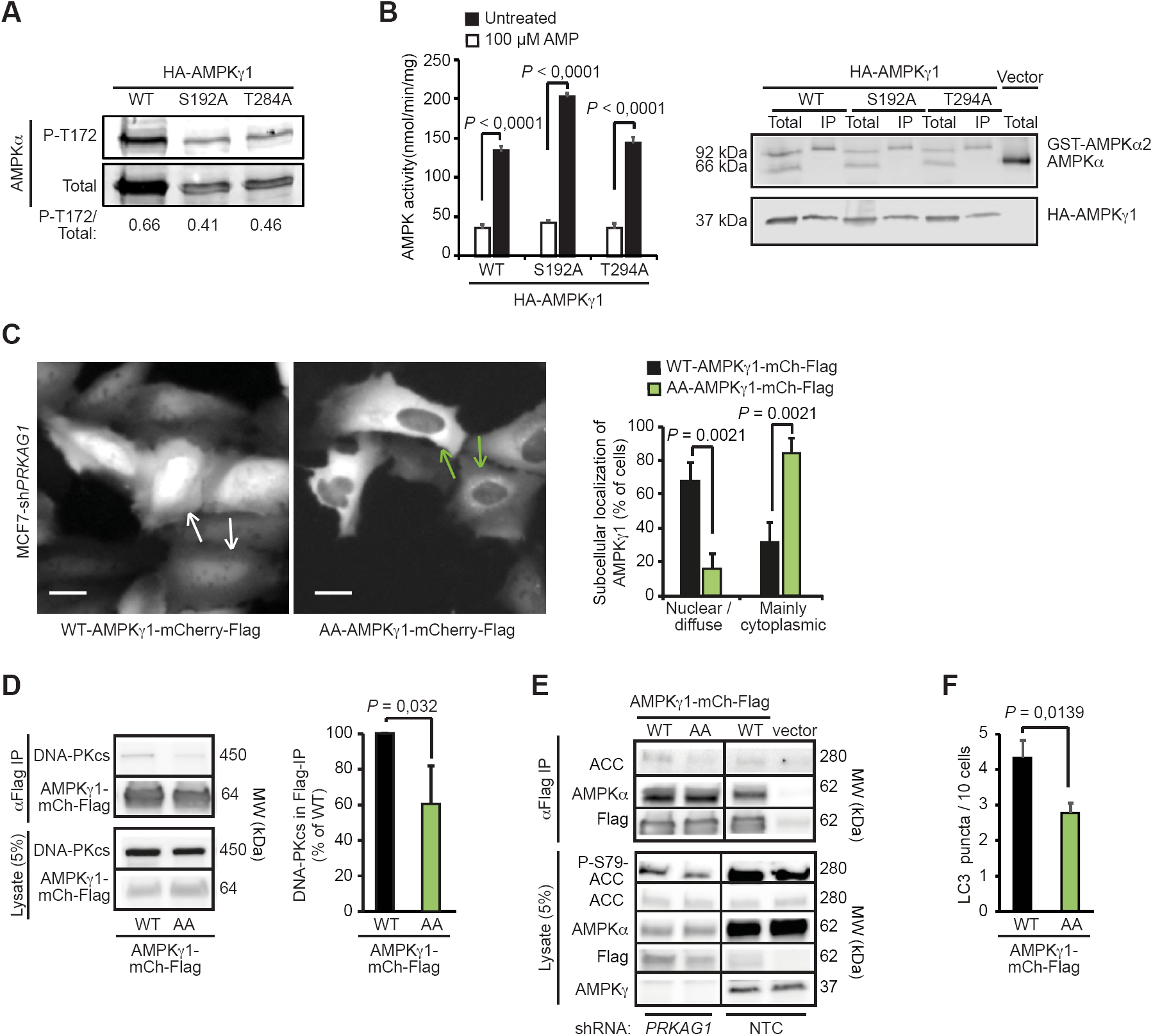
DNA-PKcs regulates AMPKγ1 localization and activity. (A)Representative immunoblots of indicated proteins from COS7 cells transfected with GST-AMPK*αα*2, AMPKβ1-Flag and either wild type (WT) HA-AMPK*γ*1 or its phosphorylation-deficient mutants (S192A or T284A). *(B)In vitro* kinase activity assay of AMPK complexes immunopurified with anti-Flag agarose gel from COS7 cells transfected as in (A) and left untreated or treated with 100 µM AMP (*left*). Values represent means of specific AMPK activity +SEM of four independent experiments. *Right*, representative immunoblots of indicated proteins from total cell lysates and anti-Flag IPs. (C)Representative images of MCF7-sh*PRKAG1* cells transiently transfected with WT or AA-mutant of AMPK*γ*1-mCh-Flag (*left*), and quantification of the subcellular localization of the transfected proteins (*right*). Values represent means +SD of three independent experiments with ≥ 50 randomly chosen cells analyzed in each sample. Arrows mark cells with nuclear/diffuse (white) and mainly cytoplasmic (green) localization of AMPK*γ*1-mCh-Flag. Scale bars, 10 µM. (D)IP of AMPK complexes with anti-Flag antibodies from lysates of MCF7 cells transiently transfected with either WT or AA mutant of AMPK*γ*1-mCh-Flag. *Left*, representative immunoblots of indicated proteins in IPs and lysates. *Right*, quantification of co-precipitating DNA-PKcs from three independent experiments +SD. (E)IP of AMPK complexes with anti-Flag antibodies from lysates of MCF7-sh*PRKAG1* and -shNTC cells transiently transfected with WT or AA mutant of AMPK*γ*1-mCh-Flag. (F)LC3 puncta in MCF7-EGFP-LC3 cells transiently transfected with WT or AA mutant of AMPK*γ*1-mCh-Flag. Values represent means +SEM of ≥ 7 samples with 10-30 cells analyzed in each. *P*-values were calculated by 2-tailed, homoscedastic student’s *t*-test.

In order to study the role of AMPK*γ*1 phosphorylation in living cells, we generated stable AMPK*γ*1-depleted MCF7 cells by lentiviral infection of *PRKAG1* shRNA (MCF7-sh*PRKAG1*). As expected, the depletion of AMPK*γ*1 resulted in the destabilization of the AMPK complex as evidenced by the disappearance of the AMPK*αα* subunit (Fig EV5A), as well as reduced growth rate and stressed appearance of the cells. Interestingly, the depletion of the AMPK complex also resulted in reduced phosphorylation of T2609-DNAPKcs (Fig EV5B), suggesting that AMPK may regulate the activation of DNA-PKcs. We then reconstituted the AMPK*γ*1-depleted cells with either wild type or a phosphorylation defective double mutant (S192A/T284A, referred to as the AA mutant) of AMPK*γ*1 fused to mCherry and Flag (AMPK*γ*1-mCh-Flag). Microscopic analysis of transiently expressed proteins revealed that contrary to the primarily nuclear localization of the wild type AMPK*γ*1-mCh-Flag, its phosphorylation defective AA mutant was predominantly in the cytoplasm of MCF7-sh*PRKAG1* cells (Fig 5C). Notably, the AA mutant formed a perinuclear rim resembling that observed for the endogenous AMPK*γ*1 upon DNA-PKcs inhibition (Figs 3D and 5C). Similar differences between the localization of wild type and AA-mutants of AMPK*γ*1-mCh-Flag were observed in U2OS cells expressing the endogenous AMPK*γ*1 (Fig EV5C). Immunoprecipitation of the AMPK complexes from transiently transfected MCF7 cells, revealed that complexes containing the AA mutant of AMPK*γ*1 had reduced affinity to DNA-PKcs and ACC and reduced capacity to phosphorylate S79-ACC, while its affinity to AMPK*αα* was unchanged (Figs 5D and 5E). In line with the reduced AMPK activity, MCF7-sh*PRKAG1* cells transfected with the AA mutant of AMPK*γ*1 had significantly fewer LC3-positive autophagic vesicles than cells transfected with the wild type protein (Fig 5F). Taken together, these data suggest that DNA-PKcs primes AMPK for the activation by upstream kinases, and that phosphorylation of S192 and/or T284 in AMPK*γ*1 is responsible for this effect.

### Phosphorylation of AMPKγ1 enhances lysosomal localization of AMPK and LKB1

Outer membranes of late endosomes and lysosomes (hereafter referred to as lysosomes) have recently emerged as important signaling platforms for major metabolic kinases, including AMPK and mTORC1 (Zhang et al., 2014). Immunocytochemistry and immunoblot analyses of purified lysosomes revealed that also a small proportion of DNA-PKcs associated with a subset of LAMP1-and ATP6V0D-positive lysosomes (Figs 6A, 6B and 6C). Therefore, we investigated whether DNA-PKcs regulated the lysosomal localization of the AMPK complex, its activating kinase LKB1 or AXIN, a scaffold protein necessary for the LKB1-mediated activation of AMPK on lysosomal membranes (Zhang et al., 2014, Zhang, Guo et al., 2013). AMPK*αα*, LKB1 and AXIN were detected in lysosomal fractions of unstimulated MCF7 cells and their lysosomal association was further increased upon short glucose starvation (Figs 6B and 6C). This increase was completely abolished upon inhibition of DNA-PKcs activity by NU7441 (Fig 6B). To test whether the glucose starvation-induced and DNA-PKcs-dependent lysosomal recruitment of AMPK and LKB1 complexes required the phosphorylation of S192 and/or T284 residues of AMPK*γ*1, we analyzed the lysosomal localization of these complexes in MCF7 cells transfected with either the wild type AMPK*γ*1-mCh-Flag or its phosphorylation defective AA-mutant. Immunoblot analyses of purified lysosomes from transfected cells revealed a reduction in the lysosomal recruitment of AMPK*γ*1 by the AA mutation both in control condition and upon glucose starvation (Fig 6C). Similar amounts of LKB1 and AXIN were found in the lysosomal fractions of unstimulated MCF7 cells expressing AMPK*γ*1-mCh-Flag constructs, but their further recruitment to lysosomes in response to glucose starvation was inhibited by the AA mutation as compared to the wild type protein (Fig 6C). The reduction in their lysosomal recruitment was associated with significantly reduced association of the AA mutant AMPK*γ*1-mCh-Flag with LKB1 (Figs 6D and EV6). The AA mutant failed also to enhance the AMPK activation upon glucose starvation as analyzed by the phosphorylation of T172-AMPK*αα* and association with ACC (Figs 6D and EV6). Based on these data, we conclude that the DNA-PKc-mediated phosphorylation of AMPK*γ*1 plays a critical role in the lysosomal activation of AMPK through the regulation of the lysosomal localization of the AMPK complex and its subsequent association with LKB1.

**Figure 6:**
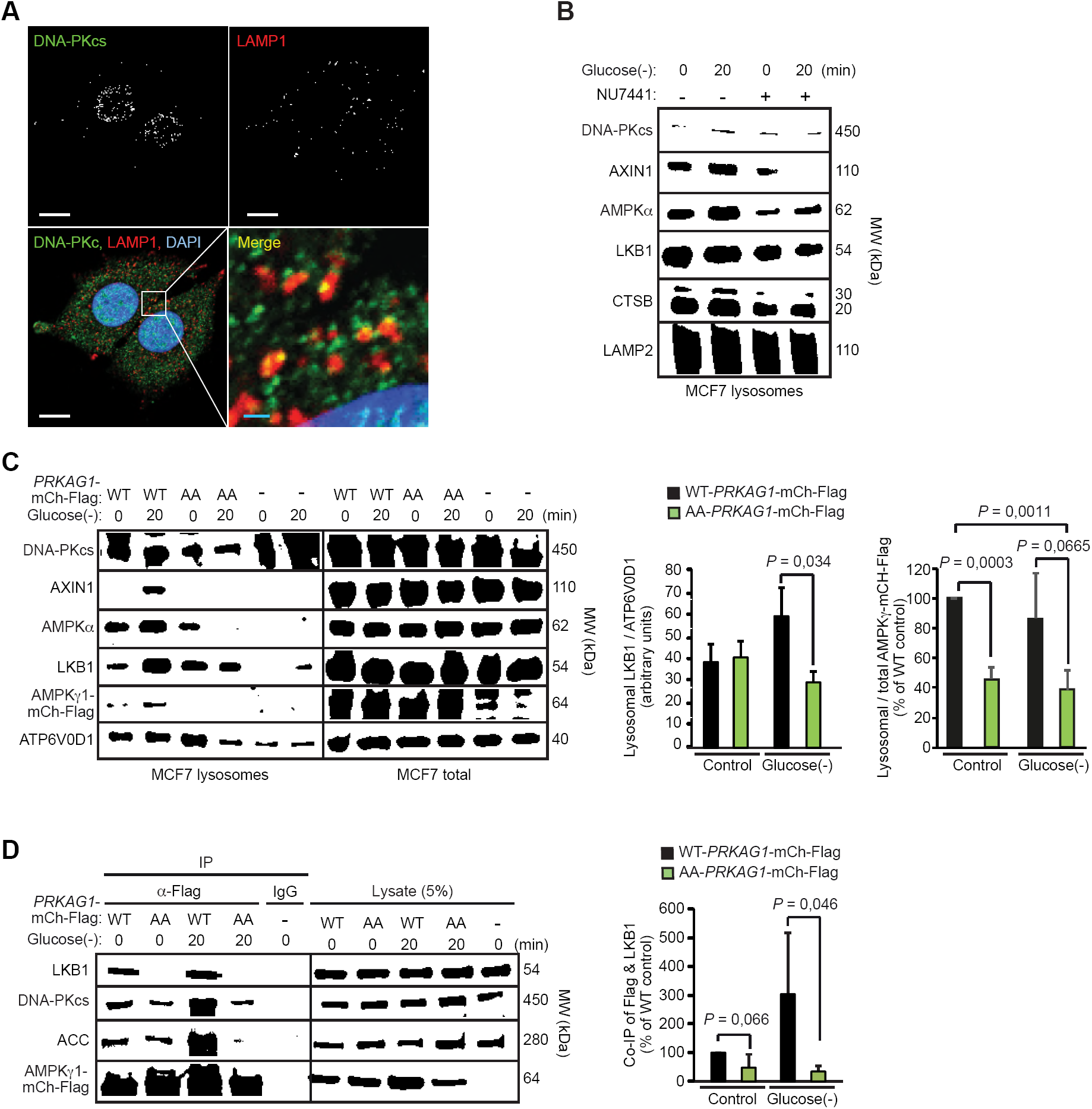
DNA-PKcs regulates lysosomal localization and LKB1 association of AMPK. (A)Representative confocal images of MCF7 cells stained for DNA-PKcs, LAMP1 and DNA (Dapi). Scale bars, 10 µM (white) and 1 µM (blue). (B)Representative immunoblots of indicated proteins from lysosomes purified from MCF7 cells starved for glucose for indicated times. When indicated (+), cells were pre-treated with 1 µM NU7441 for 16 h. (C)Representative immunoblots of indicated proteins from lysosomes and total cell lysates of MCF7 cells transiently transfected with WT or AA mutant of AMPK*γ*1-mCh-Flag or an empty vector (-) and starved for glucose for indicated times (*left*), and densitometric quantification of LKB1 / ATP6V0D1 and lysosomal / total AMPK*γ*1-mCh-Flag ratios in lysosome samples (*right*). Values represent means +SD of 3 independent experiments. (D)IP of AMPK complexes with anti-Flag antibodies from lysates of MCF7 cells transiently transfected with either WT or AA mutant of AMPK*γ*1-mCh-Flag and starved for glucose for indicated times (*left*), and densitometric quantification of LKB1 in anti-Flag IPs (*right*). Values represent means +SD of 4 independent experiments. *P*-values were calculated by student’s *t*-test.

## DISCUSSION

By exploiting RNAi technology and a luciferase-based live cell reporter assay for autophagic LC3 turnover in MCF7 breast cancer cells, we have here identified multiple genes encoding for kinases and kinase-related proteins as putative regulators of basal and etoposide-induced autophagic flux. Validating our screening approach, *bona fide* autophagy genes, phosphoinositide-3-kinase regulatory subunit 4 (*PIK3R4*) and *ULK1* (Yang & Klionsky, 2010), as well as several previously reported autophagy regulators, *e.g.* glycogen synthase kinases 3*αα* and β (*GSK3A* and *GSK3B*), inhibitor of nuclear factor kappa-B kinase subunit β (*IKBKB*) and *ATM* (Alexander, Cai et al., 2010, Criollo, Senovilla et al., 2010, Lin, Li et al., 2012), are among the candidate genes, whose depletion inhibits autophagic flux. On the other hand, the roles of the majority of the identified genes, including the three strongest candidates for regulators of either etoposide-induced or basal autophagic flux, *i.e. PRKDC*, NUAK family kinase 1 (*NUAK1*), mitogen-activated protein kinase kinase kinase 9 (*MAP3K9*), phosphoinositide 3-kinase adapter protein 1 (*PIK3AP1*), testis-specific serine kinase substrate (*TSKS*) and nuclear receptor-binding protein 2 (*NRBP2*), have not been addressed at all, or are as yet controversial. It should be noted that further validation is needed to define our candidates as true regulators of autophagy.

*PRKDC* siRNA targeting DNA-PKcs was statistically the strongest inhibitor of etoposide-induced autophagic flux and scored also as a significantly better inhibitor of basal autophagic flux than *BECN1* siRNA in our screen. Our validation enforced the requirement of DNA-PKcs for basal autophagy in MCF7, U2OS and HeLa cells as well as for that induced by etoposide, 5-FU, daunorubicin and irradiation in MCF7 and U2OS cells. Importantly, our data strongly suggest that DNA-PKcs regulates autophagy independent of its well-defined role in DNA repair as the effect of *XRCC* siRNA-mediated depletion of KU80, which is essential for DNA-PKcs-mediated DNA repair (Lees-Miller & Meek, 2003), failed to inhibit etoposide-induced autophagy, and DNA-PKcs depletion had no major effect on etoposide-induced DNA damage response as analyzed by the cell cycle profile and the phosphorylation of TP53. The role of DNA-PKcs in autophagy has been previously addressed in only few studies with somewhat controversial results (Daido et al., 2005, Zhen, Li et al., 2016). Based on the comparison of DNA-PKc expressing, radiation resistant M059K cells and DNA-PKc deficient, radiation sensitive M059J cells derived from the same malignant glioma specimen, Daido and coworkers have suggested that DNA-PKcs inhibits autophagy and protects glioma cells against radiation-induced autophagy-dependent cell death (Zhen et al., 2016). On the contrary and in line with our results, Yoon and coworkers have reported more autophagic vesicles in DNA-PKc expressing M059K cells than in DNA-PKc deficient M059K cells upon capsaicin treatment (Yoon, Ahn et al., 2012). These discrepancies could be caused by cell or stimulus specific differences. Alternatively, the observed difference in radiation-induced autophagy between M059J and M059K cells observed by Daido and coworkers could be due to other changes than DNA-PKcs expression levels. Supporting this view, M059K cells have much lower levels of ULK1 and are practically devoid of cytoplasmic ULK1 (Fig EV7), which could explain their defective radiation-induced autophagy. Further supporting the autophagy-stimulating function of DNA-PKcs, Zhen and coworkers have recently reported that pharmacological inhibitors, *PRKDC* shRNA and *PRKDC*-targeting miR-101 inhibit salinomycin-induced cytoprotective autophagy in osteosarcoma cells (Zhen et al., 2016).

The data presented above demonstrate that DNA-PKcs associates with the AMPK complex and regulates its subcellular localization and association with the major AMPK kinase, LKB1, by phosphorylating the AMPK*γ* subunit of the AMPK complex. Interestingly, these data also suggest that this regulation occurs, at least partially, on the lysosomal membrane, which is emerging as the major site for the coordination of cellular metabolism (Bar-Peled & Sabatini, 2014, Liu, Palmfeldt et al., 2018, Wunderlich, Fialka et al., 2001, Zhang et al., 2014). Supporting our findings, the association between DNA-PKcs and AMPK has been identified by two previous studies. Amatya and coworkers have demonstrated an interaction between DNA-PKcs and AMPK*γ* using yeast 2-hybrid method and co-immunoprecipitation from mammalian cells (Amatya, Kim et al., 2012), while Lu and coworkers have verified this interaction in mammalian cells by co-immunoprecipitation and immunocytochemistry (Lu, Tang et al., 2016). In line with our data, these studies have also identified DNA-PKcs as a positive regulator of AMPK activity, however, without addressing the underlying mechanism. Here, we identified S192 and T284 residues of AMPK*γ* as DNA-PKcs substrates and confirmed their necessity for DNA-PKcs-mediated activation of AMPK by site-specific mutagenesis. The AMPK complex containing the AA mutant of AMPK*γ* with both identified phosphorylation sites substituted with alanine were not only defective in stress-induced activation but showed also a dramatically altered subcellular localization, being predominantly cytoplasmic contrary to the primarily nuclear localization of the wild type protein. Supporting the role for DNA-PKcs in subcellular localization of AMPK, the localization of AMPK complex containing the AA mutant of AMPK*γ* is similar to that observed in cells where DNA-PKcs activity is inhibited by Ebstein-Barr virus latent membrane protein 1 (LMP1) (Lu et al., 2016).

Lysosomal membrane has been recently recognized as an important site for the activation of AMPK (Zhang, Hawley et al., 2017, Zhang et al., 2014). In the absence of glucose, LKB1 and scaffold protein AXIN translocate to the lysosomal membrane, where they form a large complex with AMPK, vacuolar H^+^-ATPase and Ragulator complexes (Zhang et al., 2014). This super complex, referred to as the AXIN-based AMPK activation complex (Lin & Hardie, 2018), serves as the site for LKB1-mediated phosphorylation and activation of AMPK*αα* (Zhang et al., 2014). Our data showing that both genetic and pharmacological disturbance of DNA-PKcs-mediated phosphorylation of AMPK*γ*1 inhibits the lysosomal recruitment of AMPK, LKB and AXIN strongly suggest that this phosphorylation enhances AMPK activation by promoting the formation of the AXIN-based AMPK activation complex. The constitutive nature of both the DNA-PKcs -AMPK*γ*1 association and the S192/T284-AMPK*γ*1 phosphorylation together with their decline upon AMPK activating stimuli suggest that DNA-PKcs-mediated phosphorylation of AMPK*γ*1 serves as the initiating signal for the activation of AMPK while being dispensable for the maintenance of the kinase activity. Interestingly, the N-myristoylation of AMPKβ subunits, which is necessary for the recruitment of the AMPK complex to the lysosomal membrane (Zhang et al., 2017), has been reported to serve a similar gatekeeper function, *i.e.* being necessary for the initial AMP-induced phosphorylation of AMPK*αα* but not for the subsequent allosteric activation by AMP once AMPK*αα* is phosphorylated (Oakhill, Chen et al., 2010). Thus, it is tempting to speculate that these two events are coordinated *e.g.* by DNA-PKcs-mediated phosphorylation of AMPK*γ*1 occurring only on the lysosomal membrane or AMPK*γ*1 phosphorylation being a prerequisite for AMPKβ myristoylation. In light of our present data showing that some DNA-PKcs co-localizes with lysosomes and that the lysosomal recruitment of AMPK*γ*1 is inhibited by the mutation of DNA-PKcs phosphorylation sites, both scenarios remain possible.

The ability of DNA-PKcs to regulate AMPK activation links DNA-PKcs not only to the regulation of autophagy, but also to the regulation of cellular metabolism in a more general manner. Contrary to the activating effect of DNA-PKcs on MAPK, ageing-related increase in DNA-PKcs activity has been reported to inhibit AMPK activity in aging muscle cells via phosphorylation-mediated inhibition of Hsp90 chaperone activity towards AMPK*αα*2, a muscle specific AMPK*αα* isoform (Park, Gavrilova et al., 2017). Thus, the crosstalk between DNA-PKcs and AMPK is likely to have several layers and depend on cellular context. Regarding the emerging role of DNA-PKcs in the control of cellular metabolism, it is interesting to note that DNA-PKcs itself is regulated by the metabolic state, fed conditions favoring its activation and fasting conditions inhibiting it (Wong, Chang et al., 2009). Furthermore, DNA-PKcs can regulate metabolism in an AMPK independent manner by phosphorylating the USF-1 transcription factor, which activates the transcription of fatty acid synthase and triglyceride synthesis (Wong et al., 2009).

Taken together, data presented here provide a molecular mechanism explaining how DNA-PKcs enhances AMPK activation and autophagy, thereby linking DNA damage response not only to the regulation of autophagy but to the control of cellular metabolism in general.

## MATERIALS AND METHODS

### Cell culture, reagents and treatments

The MCF7 cell line used here is the S1 subclone of the human ductal breast carcinoma cell line MCF7 selected for high TNF sensitivity (Jäättelä, Benedict et al., 1995). Human HeLa cervix carcinoma, U2OS osteosarcoma, MO59J and MO59K glioma cells as well as COS7 kidney epithelial cells were obtained from ATCC. MCF7-EGFP-LC3, -RLuc-LC3^wt^ and -RLuc-LC3^G^120^A^ and HeLa-RLuc-LC3^wt^ and -RLuc-LC3^G^120^A^ have been described previously (Farkas et al., 2009, HØyer-Hansen, Bastholm et al., 2007). U2OS-RLuc-LC3^wt^ and -RLuc-LC3^G^120^A^ were created here as described for MCF7 cells previously (Farkas et al., 2009). MCF7 cells were cultured in RPMI 1640 with Glutamax (Thermo Fisher Scientific (TFS), 61870-010) supplemented with 6% heat-inactivated fetal calf serum (TFS, 10270-106), penicillin and streptomycin (TFS, 15140-122). Other cells were cultured in DMEM (TFS, 31966-021) supplemented with 10% heat-inactivated fetal calf serum, penicillin and streptomycin. The cells were maintained in a humidified atmosphere at 37°C, 5% CO2. All cell lines were found negative for mycoplasma using Venor^®^GeM Classic PCR kit from Minerva Biolabs (11-1100). 3-methyladenine (3-MA), 5-fluorouracil, camptothecin, cisplatin, daunorubicin, doxorubicin, hydroxyurea, etoposide, and rapamycin were purchased from Sigma-Aldrich; NU7441 from Kudos Pharmaceuticals; oligofectamine from Invitrogen; and coelenterazine, enduren, and 5x passive lysis buffer from Promega. Amino acid and glucose starvations were performed in Hank’s Balanced Salt Solution (HBSS; TFS, 14025-092) and glucose free DMEM with glutamate (Gibco), respectively. Drugs were added simultaneously during co-treatments. Ionizing radiation was delivered by an X-ray generator (150 kV; 6 mA;; 0.708 Gy/min dose rate, YXLON).

### Transfections

Plasmids listed in Table EV2 were transfected to subconfluent cells with Fugene-HD (Invitrogen, E291A) and in the case of rescue experiments with Gene Juice (Novagen, 70967-3) transfection reagents according to manufacturers’ instructions. The medium was changed 16 h after the transfection and the experiments were performed 32 h later.

siRNA transfections were performed as reverse transfections using 5-20 nM siRNA and Oligofectamine transfection reagent (Promega, 12202-011) following manufacturer’s protocols. siRNA sequences are listed in Table EV2. The *Silencer*® Select Human Kinase siRNA Library V4 (Ambion, 4397918) was used in the screen.

Validated Control, *PRKAG1* and *PRKDC* shRNA lentiviral vectors (MISSION;; Sigma-Aldrich) and a lentiviral vector encoding for shRNA-resistant *PRKAG1* were amplified and infected according to standard protocols. Cells were selected with puromycin for a minimum of five passages and experiments were performed at passages 6-10.

### Site-directed mutagenesis

Site-directed mutagenesis of the constructs was carried out using QuickChange Lightning mutagenesis kit (Agilent, 210518). Oligos for mutation PCR were designed by Agilent QuickChage Primer Design web page (www.genomics.agilent.com) and are listed in Table EV2. Mutated constructs and all plasmids were confirmed by sequencing.

### Autophagic flux assay and screen

The ratio of luciferase activity between Rluc-LC3^wt^ and Rluc-LC3 ^G^120^A^ expressing cells was used as a measure for autophagic flux in living cells or cell extracts as described previously (Farkas et al., 2009). The screen was performed by measuring autophagic flux in living cells. Briefly, cells were transfected with a pool consisting of 6 nM of each of the three unique siRNAs targeting each kinase. The control siRNAs, non-targeting control siRNA and BECN1 siRNA (positive control for autophagy inhibition) were used at 18 nM. The medium was changed 52 h after the transfection with medium containing 50 nM EnduRen, and cells were allowed 8 hours to achieve equilibrium between intra-and extracellular EnduRen, before the first measurement of luciferase activity (referred to as T0) and treatment with etoposide (50 µM) or DMSO. Luciferase activity was subsequently measured 4, 8, 12 and 15 h after the treatment. Luminescence was measured using the Varioskan Flash plate-reader (Thermo Electron Corporation).

### Detection of autophagic membranes

The LC3 puncta formation in MCF7-EGFP-LC3 was assessed after fixation in 3.7% formaldehyde (Sigma-Aldrich, 252549) applying Olympus IX-70 inverted microscope with a 20x Lucullan objective with numerical aperture of 0.45 or Zeiss LSM510 microscope with a 40x Plan-APOChromat with numerical aperture of 1.3/oil. LC3-puncta formation in other cells and WIPI2 puncta formation were analyzed after methanol permeabilization and appropriate antibody staining (Table EV2).

### Immunoblotting

Immunoblotting was performed using standard protocols described previously (Corcelle-Termeau, Vindelov et al., 2016). Primary antibodies used are listed in Table EV2. The signal was detected with Clarity Western ECL Substrate (Bio-Rad, 170– 5061), and Luminescent Image Reader (LAS-1000Plus, Fujifi lm, Tokyo, Japan), and quantified by densitometry with ImageJ software (imagej.net/Fiji).

### Immunoprecipitation

Proteins from cell lysates were extracted with lysis buffer (25 mM Tris, pH 7., 150 mM NaCl, 5 mM MgCl2, 0.5% NP-40, 1 mM DTT, 5% glycerol, 1 mM phenylmethanesulfonyl fluoride, 2 μg/ml pepstatin, 2 μg/ml leupeptin, and 2 μg/ml aprotinin) in the presence of phosphatase inhibitors (Roche Applied Science, 04906837001) at 4 °C. Cellular debris was removed by centrifugation at 13,000 ×g for 10 min at 4 °C. One milligram of protein from the cell lysates was immunoprecipitated for 4-16 at 4°C, using indicated antibodies listed in Table EV2 and protein G (Sigma-Aldrich, P3296) or protein A (GE Healthcare, 17-5138-01) agarose beads. After the last washing step, SDS-PAGE buffer was added (40 µl) and samples were heated at 95°C for 10 min. 15 – 30 μl of immunopurified proteins/lane were separated on a 4 – 15 % polyacrylamide, precast Mini-PROTEAN TGS gel (BIO-RAD) followed by transfer on Trans-Blot Turbo Transfec Pack (0.2μm) nitrocellulose (BIO-RAD), and immunoblotting with indicated antibodies listed in Table EV2.

### Immunocytochemistry and confocal microscopy

Cells grown on coverslips and fixed either in methanol (-20 C^^0^^) for 2 min or in 4% PFA in PBS for 10 min were washed with PBS, quenched with 50 mM NH_4_Cl for 10 min, washed with PBS and blocked in buffer 1 (1% BSA, 0.3% TritonX-100, 5% goat serum in PBS) for 20 min before incubation with indicated primary antibodies (Table EV2) in buffer 1 for 16 h at 4 C°. After three washes in buffer 3 (PBS, 0,05% Tween-20), coverslips were incubated with appropriate secondary antibodies (Alexa Fluor® 488-conjgated donkey anti-mouse IgG or Alexa Fluor® 568-conjugated donkey anti-rabbit IgG from Promega (A-21202, A-10042)) for 1 hour in buffer 2 (0,25% BSA, 0,1% TritonX-100, PBS), washed three times in buffer 2, mounted with Prolong antifade Gold with DAPI (Life Technologies, P36935) and dried for 24 h at 20° C.

### Generation of Antibodies

Phosphorylation specific (P-S192 and P-T284) polyclonal AMPKγ1 antibodies were produced in rabbit using a peptide synthesized by JPT Peptide Technologies GmbH (Germany) with sequences listed in Table EV2. Prior to immunization peptides were coupled to Imject maleimide-activated Keyhole Limpet Hemocyanin (mcKLH, Pierce 77605) by GenScript (www.genscript.com) according to manufacturer’s protocol. Antibodies were purified from positive sera using peptide antigen affinity columns prepared with a standard protocol (sulfo-link immobilization columns, Pierce 44999) with immobilized peptides against phosphorylated peptides following the manufacturer’s protocol. After low pH elution (50 mM glycine [pH 2.5]), the antibodies were cleared for cross-reactivity on columns with synthetic peptides corresponding to the non-phosphorylated forms of S192 and T284 peptides immobilized to peptide antigen affinity columns and eluted with low pH (50 mM glycine (pH 2.5). Antibodies were stored as aliquots in PBS, 50% glycerol, 5 % BSA, and sodium azide at –20° C. The specificity of antibodies was tested against phosphorylated peptides immobilized to nitrocellulose membranes (Bio-Rad) by dot blotting.

### Protein purification and *in vitro* kinase activity assay

AMPK*γ*1-mCherry-Flag proteins were purified from transiently transfected MFC7 cells with Anti--FLAG® M2 Affinity Gel (Sigma Aldrich, A-2220) using RIPA lysis buffer (50 mM Tris (pH 7.5), 150 mM NaCl, 50 mM NaF, 1 mM EDTA, 1 mM EGTA, 1% TritonX-100, 0,5 % deoxycholate, 0,05% SDS). The immune complexes were washed extensively with RIPA buffer and three times with DNA-PK reaction buffer (50 mM HEPES (pH 7.5), 100 mM KCl, 10 mM MgCl^^2^^, 0.2 mM EGTA, 0,1 mM EDTA, 1 mM DTT, 5 μg Calf thymys DNA). For the in vitro kinase reaction, the FLAG-purified proteins were either immobilized to FLAG-beads or released before kinase reaction by using excess amount of FLAG-peptide.

For the DNA-PK kinase assay, indicated AMPK*γ*-Cherry-FLAG proteins were incubated at 33° C for 20 min in DNA-PK reaction buffer supplemented with 0.2 mM ATP, 0.2 μg/ml BSA, 0.2 µl *αα*-^^32^^P -ATP (3000 Ci/mmol) and 250 U purified DNA-PK (Promega, V581A). Reaction was terminated by adding SDS sample buffer and subjected to SDS–PAGE following the blotting of the SDS-PAGE samples on to nitrocellulose filters. Incorporation of AMPK*γ* labeled phosphate was quantitated using an autoradiography with phosphoimager plates following plate read by Typhoon 9410 variable mode imager (Amersham Biosciences). Typhoon 9410 raw files were further quantitated by MultiGauge V3.0 (FujiFilm).

To test the DNA-PK-mediated phosphorylation of peptides, 2 µg of non-phosphorylated (S192: amine-PEFMSKSLEELQIGC-amide;; T284: amine-KCYLHETLETIINRLC-amide) or phosphorylated (P-S192: amine-PEFMSK-pS-LEELQIGC-amide;; P-T284: amine-KCYLHE-pT-LETIINRLC-amide) AMPKγ1 peptides (JPT peptide technolgies GmbH) corresponding to predicted DNA-PK phosphorylation sites were diluted in DNA-PK reaction buffer supplemented with 1 µM ATP, 0.2 μg/ml BSA, 10 µCi *γ*-^^32^^P -ATP (3000 Ci/mmol) and 250 U purified DNA-PK and incubated at 34°C for 20 min, heated to 60 °C and spotted to nitrocellulose membrane. Dried peptide containing filters were fixed with glutaraldehyde (3%) in PBS at at 20° C for 5 min, washed 3 x 5 min with PBS under vigorous agitation before the incorporation of labeled phosphate to the peptides was quantified as above.

### *In vitro* AMPK activity assay

Heterotrimeric AMPK complexes consisting of GST-AMPK*αα*2, AMPK1-FLAG and wild type or mutated HA-AMPK*γ*1 were immunoprecipitated from COS7 cells transfected with corresponding plasmids (Table EV2) using Anti-FLAG M2 affinity agarose gel and washed repeatedly with buffer A (50mM HEPES pH 7.4, 150mM NaCl, 10% glycerol and 0.1% Tween-20) prior to activity assay. AMPK activity assays using radiolabeled [*γ*-^^32^^P]-ATP were conducted as described previously (Ngoei, Langendorf et al., 2018, Oakhill, Scott et al., 2018).

Assays were conducted in the presence of 100 µM SAMS peptide (NH2-HMRSAMSGLHLVKRR-COOH), 5 mM MgCl2, 200 µM [*γ*-^^32^^P]-ATP for 10min at 30°C in the presence/absence of 100 µM AMP. Phosphotransferase activity was quenched by spotting onto P81 phosphocellulose paper (Whatman, GE Healthcare) followed by repeated washes in 1% phosphoric acid. ^^32^^P-transfer to the SAMS peptide was quantified by liquid scintillation counting (Perkin Elmer).

### Purification of lysosomes

Subconfluent cells grown in 15 cm petri dishes in 20 mL complete medium were treated for 18 h with 0.2 mL iron dextran solution (53 mg/mL in deionized water;; prepared essentially as described previously (Diettrich, Mills et al., 1998)), washed in medium without iron dextran, and cultured for additional 4 h in complete medium before the indicated treatments. After treatments, cells were scraped off in ice cold KPBS (136 mM KCl, 10 mM KH2PO4, pH 7.25 Adjusted with KOH) and centrifuged at 1000x g for 2 minutes at 4° C. The pellet was dissolved in 950 µl of KPBS and homogenized with 10 strokes through a 25-gauge needle, and centrifuged at 1000 g for 2 min at 4° C. The supernatant was loaded to prewashed 3 ml MACS –magnetic columns prewashed with KPBS and placed into MiniMacs magnetic separator at 4° C. Loaded columns were washed three times with 1 ml of KPBS and bound lysosomes were released from the matrix by removing the MACS-column from the magnetic field and applying 80 µL of KPBS buffer on the column. The eluted material was mixed with 80 µl of 2x SDS-PAGE buffer and analyzed for immunoblotting.

### Cell cycle analysis

Cells were treated as indicated, fixed in methanol, and stained with propidium iodine. Flow cytometry (FACSCalibur, BD Biosciences) was performed on 10,000 cells/sample and data were analyzed using the FlowJo software (www.flowjo.com).

### Statistical analyses

For statistical analysis of the etoposide screen, we applied a linear regression model to study inhibition over time. siRNAs that inhibited etoposide-induced autophagic flux significantly better than *BECN1* siRNA were identified by comparing slopes (inhibition per time unit) and tested by means of Wald tests. Differences in slopes that yielded a p-value < 0.05 were considered statistically significant. Other P-values were calculated by 2-tailed, homoscedastic student’s *t*-test.

## ACKNOWLEDGMENTS

We thank Developmental Studies Hybridoma Bank developed under auspices of the National Institute of Child Health and Human Resources and maintained by The University of Iowa as well as Jiri Lukas and Jiri Bartek for valuable research reagents.

## Funding

M.J. was supported by the Danish National Research Foundation (DNRF125), European Research Council (AdG 340751), Danish Cancer Society (R40-A1793), Danish Council for Independent Research (0602-02707B) and Novo Nordisk Foundation (12OC0001341). J.S.O was supported by grants from the National Health and Medical Research Council (NHMRC) and the Australian Research Council (ARC), and in part by the Victorian Government’s Operational Infrastructure Support Program. We would also like to acknowledge the support of the TRANSAUTOPHAGY COST Action (CA15138).

## AUTHOR CONTRIBUTIONS

A.K. designed and performed the kinome screen and the majority of the experiments validating DNA-PKcs as an autophagy regulator. P.P. identified AMPK*γ*1 as a DNA-PKcs substrate and designed, performed the majority of the experiments validating this finding and participated in writing the first draft of the manuscript. K.N. and J.O. designed and performed the experiments addressing the ability of AMPK to bind AMP. E.C.-T. assisted in the immunocytochemistry and autophagy assays. S.P.L. and T.F. assisted in the screen and identification of the DNA-PKcs substrates. K.K.A. performed statistical analyses related to the screen. M.J. designed the overall study, supervised the experiments, assisted in the data analyses and wrote the final draft of the manuscript. All authors contributed to the final text and approved it.

## DECLARATION OF INTERESTS

Authors declare no competing interests.

## EXTENDED VIEW

Extended view includes two table (Tables EV1 and EV2) and seven figures (Figs EV1-EV7).

